# Measurements of Subthreshold and Fast Population Neuronal Activity with Genetically Encoded Calcium Indicator GCaMP-6f

**DOI:** 10.1101/505834

**Authors:** Pinggan Li, Xinling Geng, Huiyi Jiang, Adam Caccavano, Stefano Vicini, Jian-young Wu

**Affiliations:** Department of Pediatric Neurology, Sun Yat-Sen Memorial Hospital, Sun Yat-Sen University, Guangzhou, China; Department of Neuroscience, Georgetown University, Washington DC, United States; School of Biomedical Engineering, Capital Medical University, Beijing, China; Department of Pediatrics, The First Hospital of Jilin University, Changchun, China; Department of Pharmacology and Physiology, Georgetown University, Washington DC, United States

**Author notes:** These authors contribute equally (Pinggan Li, Xinling Geng).

**Keywords:** GCaMP, calcium signals, hippocampal slice, voltage-sensitive dye imaging, local field potential recordings, theta oscillation, hippocampal sharp wave

## Abstract

GCaMP-6f is among the best calcium indicators and has been widely used for monitoring neuronal activity in the brain. Applications are at cellular level (calcium transients of action potentials) or population activity (fluorimetry) during network events. Two important issues remain less explored: 1) Is GCaMP-6f signal sensitive enough for detecting subthreshold activity, similar to the sensitivity of local field potential (LFP)? 2) Is the GCaMP-6f signal fast enough for detecting network oscillations seen in LFP? Here the two issues are explored in a number of network events including hippocampus sharp waves (SWs), carbachol induced theta oscillations, interictal-like spikes and neuronal response evoked by high frequency stimuli. SWs are a typical network event with the majority of neurons receiving subthreshold excitatory or inhibitory synaptic input without firing action potentials. The excitatory/inhibitory post synaptic potentials (EPSP/IPSP) in the neuropil become detectable in local field potential (LFP) signals. We compare simultaneously recorded LFP and optical recording of GCaMP-6f fluorescent signals in Thy1-GCaMP-6f mice hippocampal slices. We found that the occurrence of SWs produces a clear population GCaMP-6f signal of 0.3% dF/F. This population GCaMP-6f signal correlated well with the LFP, albeit a delay of ~50 ms was observed. The population GCaMP-6f signal follows well with the 20 Hz population activity evoked by electric stimuli, while activity up to 40 Hz was detected with reduced amplitude. GCaMP-6f and LFP signals showed a large amplitude discrepancy. The amplitude of GCaMP increased ~1000 times from SW to carbachol induced theta burst, while the LFP changed less than 10 times. Our results suggested that population GCaMP-6f signals may become a sensitive tool for detecting network activity, especially for that with low LFP amplitude during elevated spiking rate but asynchronized events. GCaMP signal is fast enough for monitoring theta and beta oscillations (<25Hz) in the neuronal population. Faster calcium indicators (e.g., GCaMP-7) may improve the frequency response property for detecting gamma band oscillations. In addition, population GCaMP recordings are non-contact and free from stimulation artifacts. These features may be useful for high throughput recordings and applications sensitive to stimulus artifact, e.g., monitoring response during continuous stimulations.

## 1 Introduction

Calcium fluorescent signals have been used and are becoming widely used for monitoring neuronal activity in the brain thanks to the availability of genetic encoded indicators. GCaMP-6f is among the best calcium indicators to date, with high sensitivity, high fluorescent yield and relatively fast response time (Chen et al., 2013). There are two major applications of GCaMP-6f: To visualize somatic calcium transient (‘blinking’ neurons when they fire action potentials) [e.g., (Svoboda et al., 1997; Chen et al., 2013; Muto et al., 2013; Dana et al., 2014)] and monitoring population neuronal activity, (aka calcium fluorometry e.g., (Cui et al., 2013; Kupferschmidt et al., 2017)). Two important issues related to the latter remains less explored: 1) Is GCaMP-6f signal sensitive enough for detecting subthreshold activity? 2) Is the GCaMP-6f signal fast enough for detecting network oscillations?

For the sensitivity issue, we ask whether population GCaMP signal is comparable to Local field potential (LFP) recordings, i.e., is capable to detect most of the network activity in which only a small fraction of neurons fires action potentials, while the majority of neurons only have subthreshold potentials. The answer appears to be a No for two reasons: A) Fluorescence signal appears to reflect suprathreshold calcium influx, while the source of LFP can be from subthreshold synaptic currents. B) The source of the GCaMP signal appears to be mainly somatic calcium transients, while the source of the LFP voltage appears to be dendritic currents. However, measuring population subthreshold calcium signals might become possible when the signals from a large number of dendrites are integrated in the neuropil, given that the sensitivity and fluorescent yield of GCaMP-6 is excellent.

Concerning the frequency response issue, the general consensus is that the time course of somatic calcium transient is about 1 second, which is too slow for detecting most network oscillations [e.g., in (Xing and Wu, 2018), 2 Hz signals were detectable but not 10 Hz signals]. However, the rising time of calcium events is very fast and may be able to follow fast oscillations. Measurements with organic calcium indicator magnesium green showed <1 ms rising time (Regehr, 2000). When action potentials occur, the duration of intracellular calcium transient may be long and limited by the ability of intracellular buffering and clearance(Helmchen and Tank, 2015; Neher, 2013). In contrast, during subthreshold events with low calcium influx, the rising time of calcium signal would be limited by only by the onset time of the channels and the rate of calcium influx. The estimated time delay for the opening of calcium channels is only a few milliseconds (Demuro and Parker, 2004; Shuai and Parker, 2005). During subthreshold events the rate of calcium influx is related to the opening probability of low voltage activated channels (LVAs) [Reviewed by (Catterall, 2000; Grienberger and Konnerth, 2012). The opening probability of the LVA channels should be correlated to the fluctuation of membrane potential. In theory, recording fast oscillations in neuronal populations with calcium indicator is possible when large fraction of neurons has subthreshold membrane potential oscillations, and the ability of intracellular calcium buffering/clearance is much larger than the calcium influx rate. GCaMP-6f has a signal onset delay of ~40 ms (Chen et al., 2013), and it should allow up to 25 Hz of the oscillations to be detected.

Hippocampus sharp waves (SWs) are a typical network events with a small fraction of neurons firing action potentials (Ylinen et al., 1995; Csicsvari et al., 1999a; b; Mizuseki and Buzsaki, 2013) while the majority of neurons receiving subthreshold excitatory or inhibitory synaptic inputs(Hajos et al., 2013). The population summation of the EPSPs/IPSPs generates a clear voltage source/sink pair in LFP recordings [(Maier et al., 2009), reviewed by (Buzsaki, 2015). In this report we use SWs as a population neuronal event for testing if subthreshold event can be seen in the GCaMP signals. We speculate that the population summation of the calcium influx during subthreshold events would become detectable in GCaMP-6f optical recording.

We compared simultaneously recorded LFP and optical recording of GCaMP-6f fluorescent signals in Thy1-GCaMP-6f mouse hippocampal slices during SWs, interictal spikes and carbachol induced theta oscillations. Population activation by electric shocks was also used to test the frequency response characteristics of GCaMP population signals.

We found that SWs can be clearly detected optically in the population GCaMP-6f signals, and the GCaMP optical signals correlated well with the LFP, albeit a delay of ~50 ms was observed. The population GCaMP-6f signals follow well with the network activity below 20 Hz, while activity up to 40 Hz were detected with reduced amplitude. GCaMP-6f and LFP signals showed a large amplitude discrepancy. The amplitude of GCaMP increased ~1000 times from SW to carbarchol induced theta burst, while the LFP changed less than 10 times.

Our results suggested that population GCaMP-6f signals have a sensitivity comparable to that of LFPs, and may be even more sensitive than LFP for detecting network events with low LFP amplitude but unproportionally large GCaMP-6f signals. We also found that the population GCaMP signals are fast enough for monitoring theta (4-7 Hz) and beta (14-30Hz) oscillations in the slice, and the detecting limit can be pushed up to 40 Hz. Our results suggest that optical recording of population GCaMP signals may become a complementary method for the LFP. Our results also have implication for the interpretation of data in vivo obtained with the increasingly widespread use of GCaMP based photometry (Cui et al., 2013; Kupferschmidt et al., 2017).

## 2 Methods

### 2.1 Slice reparation

P21-P33 male and female C57BL/6J-Tg (Thy1-GCaMP-6f) GP5.5Dkim/J. mice (Jax 024276) mice were used to prepare paired hippocampal hemi-slices in accordance with a protocol approved by the Institutional Animal Care and Use Committee at Georgetown University Medical Center. Following deep isoflurane anesthesia, animals were rapidly decapitated. The whole brain was subsequently removed and chilled in cold (0° C) sucrose-based artificial cerebrospinal fluid (sACSF) containing (in mM) 252 sucrose; 3 KCl; 2 CaCl2; 2 MgSO4; 1.25 NaH2PO4; 26 NaHCO3; 10 dextrose; bubbled with 95% O2, 5% CO2. Hippocampal slices (480 μm thick) were cut in horizontal sections from dorsal to ventral brain with a vibratome (Leica, VT1000S). Slices were incubated in ACSF for at least 2 hours before each experiment. ACSF used for maintenance and recording contained (in mM) 132 NaCl; 3 KCl; 2 CaCl2; 2 MgSO4; 1.25 NaH2PO4; 26 NaHCO3; 10 dextrose; bubbled with 95% O2, 5% CO2 at 26° C.

### 2.2 Local Field potential (LFP) Recording

LFP recordings were done in a submerged chamber, and slices were placed on a mesh that allowed perfusion on both sides at a high flow rate (10-30 ml /min) (Hajos and Mody, 2009; Maier et al., 2009). We use low resistance glass microelectrodes (~150kΩ tip resistance). The electrodes were pulled with a Sutter P87 puller with 6 controlled pulls and filled with 0.5 M NaCl in 1% agar, which prevents leakage of the electrode solution that could potentially alter the tissue surrounding the electrode tip. The recording electrode was placed in CA1 stratum pyramidale where sharp waves have large amplitudes (Maier et al., 2009) in healthy slices.

### 2.3 GCaMP Fluorescent Recording

The GCaMP signals were recorded by a 464-channel photodiode array (WuTech Instruments). The two-stage amplifier circuits in the diode array subtract the resting light intensity and amplify the small optical signals 100 times before digitization. This achieves a 21-bit effective dynamic range to fully digitize a signal of ~0.5% dF/F. (for a recent review of the two-stage imaging system, see (Liang et al., 2015))

The dF/F is defined as (X0-F0)/F0, where X0 is the signal trace from each detector and F0 is the baseline fluorescent intensity. The signals were digitized at 1,616 frames/sec. The LFP and stimulation signals were sampled and digitized concurrently with the VSD signals. Optical imaging was performed on an upright microscope (Olympus BX51 WI) with an epi-illumination arrangement: excitation light (a 470 nm LED, ThorLabs) passes a GFP filter cube (Chroma, excitation 425-475nm, dichroic mirror 480 and emission filter 485 long pass). The GCaMP signals were imaged at two spatial resolutions: With a 20X objective (0.95 NA, Olympus) which can get blinking cells and population signals from the same tissue, and a 10x objective (0.30 NA, AMscopes) which allowed for all hippocampal subfields in the same field of view. The aperture of the diode array was 19 mm in diameter; containing a hexagonal arrangement of 469 optic fibers. The diameter of the fiber was 750 μm. Each detector on the array (pixel) collected florescent signals from an area of 37.5 μm in diameter with the 20X objective (0.95 NA, Olympus), and about 75 μm in diameter with the 10 X objective (0.4 NA American Optics). The total beam power of the LED was ~350mW at 1A. The fluorescent intensity on each detector was about 20,000 photoelectrons/ms (Illumination intensity at the sample was < 20mV/mm) when 20X 0.95NA objective was used.

### 2.4 Voltage-Densitive Dye (VSD) Imaging

VSD imaging was used to validate the linearity of the GCaMP signal (Figure 3). In 3 slices (n=3) VSD and GCaMP signals are from the same tissue and imaged by the same diode array. The slices are stained by an oxonol dye, NK3630 (Nippon Kankoh-Shikiso Kenkyusho Co., Ltd., Japan), as an indicator of transmembrane potential. The slice was stained with 5 - 10 μg/ml of the dye dissolved in ACSF for 120 minutes (26°C). During staining, the ACSF was circulated and bubbled with 95% O2 - 5% CO2. After staining, the slices were transferred back to the incubation chamber for at least 1 hour before each experiment. NK3630 binds to the external surface of the membrane of all cells and report their membrane potential change (for a recent review of the diode array and NK3630, see (Liang et al., 2015)). The absorption spectrum of the dye shifts linearly with the changes in the membrane potential (Ross et al., 1977). The VSD signal in this report is the change in absorption of light with a 705 nm wavelength. In all experiments, the detectable signals are a change in light intensity that is roughly 0.01% to 0.1% of the resting light intensity. Staining with this dye does not cause noticeable changes in spontaneous or evoked neuronal activity (Jin et al., 2002; Huang et al., 2004), and stained slices maintain viability for up to 24 hours. In 705 nm recording light, NK3630 molecules do not generate fluorescence, so no noticeable phototoxicity is detected (Jin et al., 2002).

The VSD signals were recorded by the same diode array. With a transillumination arrangement, neurons through the whole thickness of the slice (490 μm) contribute relatively equally to the VSD signal. A tungsten filament lamp was used for illumination and a 705/10nm interference filter (Chroma) was placed in the illumination path during optical recording.

During imaging experiments, the slice was continuously perfused in a submersion chamber with ACSF (same as the incubation solution) at 26°C and at a rate of more than 20 mL/min. Intermittent imaging trials were performed, with 2-3 min intervals between trials. The total light exposure for each slice was less than 600 sec, far below the dose for detectable dye bleaching or phototoxicity.

### 2.5 Stimulation

Stimulation to the CA3 area was provided with a concentric metal electrode (FHC CBDSE 75). Stimulation pulse was 0.1ms wide generated by a Master 8 stimulator (AMPI). The stimulation current was 20-100μA generated by an isolator (AMPI).

### 2.6 Data Analysis

SW events were identified by the threshold of the signal. The raw LFP traces were digitally filtered between 1-30 Hz, with a threshold set manually above the baseline noise to identify the majority of SW events. Custom programs were written in Labview for digital filtering, threshold detection, and determining the amplitude and frequency distributions. Further analysis to determine stimulus success/failure rate and latency was conducted with custom programs written in Matlab. The peak amplitude of SW events was measured from the baseline to the peak of the 1-30 Hz filtered LFP. The root mean square power (RMS) of the SW was calculated as the RMS of the GCaMP signal in a 60 ms window starting from the SW peak in the LFP signals.

Statistics were conducted in Graphpad Prism 7.0. To compare differences in mean we performed 1way ANOVA with Tukey multiple comparisons correction. All error bars displayed are SEM. * p < 0.05, ** as p < 0.005.

## 3 Results

### 3.1 Spontaneous SW Detected by Population GCaMP Signals

Spontaneous SWRs reliably occur in most hippocampal slices as reported by our previous papers and other groups (Kubota et al., 2003; Maier et al., 2003; Colgin et al., 2004; Behrens et al., 2005; Jiang et al., 2018; Sun et al., 2018). Our first goal was to test if GCaMP-6f signals can detect spontaneous SWR in hippocampal slices from the Thy1-GCaMP-6f mice. One-to-one correlations was found between LFP and GCaMP signals during spontaneous SW. The GCaMP signals were clearly seen in single trials, with a dF/F of 0.1 - 1.0% (Figure 1 A, B). Different from the localized cellular calcium transient (Miyawaki et al., 2014), the population SW signals were reliably seen a large area of the hippocampal tissue (Figure 1 B). Under a 20X objective, each of optical detector receive light from an area of 37.5 μm in diameter, so that the population signals we referred to, in this report, is a summation of dF/F from both soma and dendritic tree over a large number of neurons in the neuropil under each optical detector.

**Figure 1.**
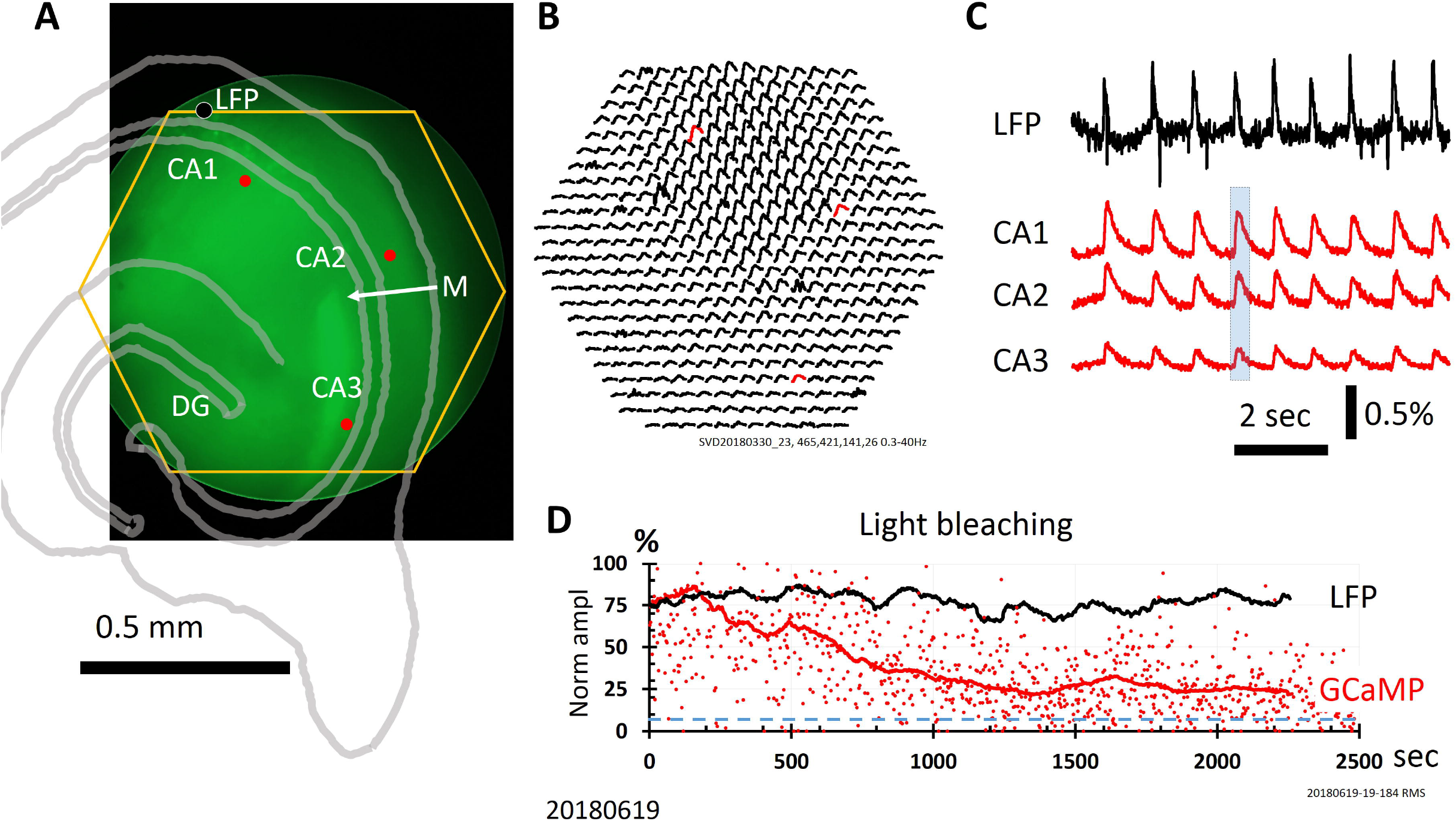
GCaMP signals of SW over hippocampus subfield. **A.** Fluorescent of GCaMP of the slice. The end of Mossy fibers (M) was used for identifying CA3, CA2 and CA1 areas. Yellow hexagon marked the field of view of the diode array. The black dot near the top left corner of the diode array mark the location of the LFP electrode (LFP). These structures were also outlined with an image of transmitted light (gray lines). **B.** dF/F from all 464 diodes during a SW. This SW event was one of the nine SWs occurred during a recording trial (marked by blue box in C). Note that GCaMP signals of SW were seen in a large area over CA1, CA2 and CA2 areas. **C.** Signals from three detectors over the CA1, CA2 and CA3 areas (Marked as red dot/trace in A and B) were plotted together with the LFP recording. The GCaMP signals of SW was about 0.5% dF/F, and the signal-to-noise ratio was high. **D.** Declining of optical signals over long recording time. Red dots mark the relative amplitude of individual SW and the red/black lines are the averages 100 SWs in a sliding window. Blue broken line marked the level of noise. Note that the LFP amplitude did not declining over time, suggesting that the declining is caused by bleaching of the GCaMP proteins.

The end of suprapyramidal mossy fibers marks the boundary between CA3 and CA2 (Blackstad et al., 1970; Gaarskjaer, 1978). In the Thy1-GCaMP-6f mouse the mossy fibers showed bright green fluorescence. The end of the mossy fiber bundle was used as a landmark of the end of CA3 area (Figure 1A, M) and CA2 area was darker as devoid of fluorescent cell bodies. CA3, CA2 and CA1 areas were identified using this landmark. When the end of the mossy fiber bundle was positioned at the center of the field of view, the 464 detectors on the diode array (yellow hexagon in Figure 1) covered a large area, including CA3, CA2 and CA1. Figure 1B showed the GCaMP signals of a single SW event from all 464 optical detectors. Signals from three detectors in the CA1, CA2 and CA3 areas were displayed with the LFP signals (Figure 1C, light blue box marked the SW event displayed in Figure 1B). In this 9-second recording trial there were 9 SWs; all show peaks in the population GCaMP signals (red traces). The signal-to-noise ratio was about 10, allowing clearly distinguishable from the noise. LFP signals were simultaneously recorded with the GCaMP signals (both sampled at 1616Hz). From 11 slices we have recorded ~6,500 SW events optically, all with a clear one-to-one correspondence between LFP and optical recording of the GCaMP signals.

### 3.2 Signal Polarity in Hippocampus Laminar

A marked difference between LFP and GCaMP signals was the signal polarity in different laminar area. LFP signals from the str. Oriens and str. Radiatum have an opposite polarity (Maier et al., 2009), known as a current source-sink pair around the stratum pyramidale (Ross et al., 1977; Johsdon and Wu, 1994). The polarity reversal was not seen in the GCaMP signals (Figure 1B). The GCaMP signals from soma (stratum pyramidale) and neuropil (polymorphic or molecular layers) had the same polarity (increased dF/F at SWS onset), suggesting that the calcium signals were not related to current flow direction in the population.

### 3.3 Recording Time

Exposure to the recording causes bleaching to the GCaMP fluorescent protein. The SW signal amplitude reduces with the exposure time. Figure 1D showed a photobleaching experiment with continuous optical recording of 2,500 seconds. In this experiment the excitation light intensity was reduced to 1/2. Under this light intensity the root mean square power (RMS) of GCaMP peak was about 5 time of the baseline RMS noise. After ~1000 second of exposure, the SW optical signal reduced to one-half of the power (Figure 1D), but it could be distinguished after 2500 seconds of exposure. Similar long-time recording was done in five slices from 5 animals, and all showed reliable recording within 1500 seconds of recording with continuous illumination. In two animals we tested long time optical recording, the optical signals remained detectable after 7200 seconds of light exposure.

### 3.4 Time Delay Between the GCaMP and VSD Signals

GCaMP signal showed a significant delay compared to local field potentials (Figure 2). The delay was about 50 ms (Figure 2C), and can be clearly seen in trace recordings (Figure 2 A, B, red traces). The delay time was verified with voltage-sensitive dye recording. In three animals we stained the slices with voltage sensitive dye NK3630, and the dye related absorption signal was measured at 705 nm. The 705 nm light did not excite the GCaMP protein but absorbed by the voltage dye bound to the neuronal membrane. The voltage signal (absorption of the NK3630) showed a delay less than 5 ms, demonstrating a good correlation between the population summation of membrane potential fluctuation and LFP signal during SW events. The delay of the population GCaMP-6f signal was comparable to the intracellular measurement of 40 ms rising time (Chen et al., 2013).

**Figure 2.**
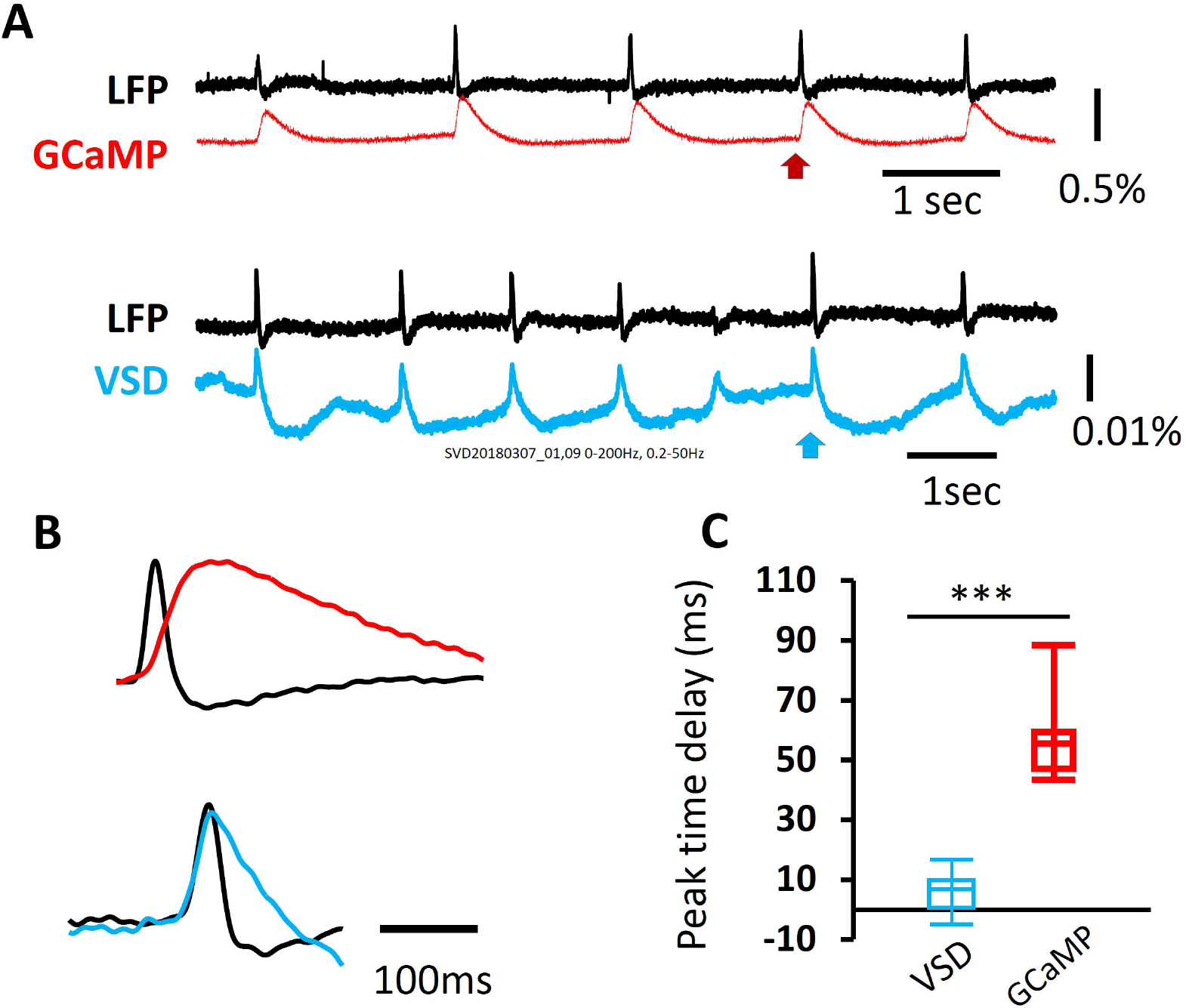
Delay in onset time. **A.** GCaMP, VSD and LFP recordings (blue, red and black traces) of SWs in the same tissue. Note that the GCaMP signal was about 50 times larger and slower. Red and blue arrow mark the SWs displayed in B. **B.** In expanded time scale, GCaMP signal (red) show marked delay compare to LFP. Such delay was not seen in the VSD signals from the same tissue (blue). **C.** Box and whiskers display of LFP-GCaMP(red) and LFP-VSD (blue) peak time delay from 26 SWs in VSD and 46 SWs, *** P<0.005.

### 3.5 Frequency Response Property

The time delay of ~50 ms of the population GCaMP-6f optical signal should in theory allow following up to 20 Hz of network oscillations in the tissue. It may also be possible to detect higher frequency signals at attenuated amplitude. To test the frequency limit of GCaMP-6f population signals, we used electric stimulation to CA3 and measured the evoked population response in the CA1 area. The stimulation intensity was low, adjusted so that the evoked response had a similar amplitude of spontaneous SWs in the same tissue (Figure 3A). Evoked GCaMP signals were also seen in the CA3 and CA1 areas (red and orange traces in Figure 3A). When two stimuli were delivered close in time, the response to the second stimulus was larger (Figure 3A, arrowhead), suggesting pair pulse facilitations was detected by population GCaMP signals.

**Figure 3.**
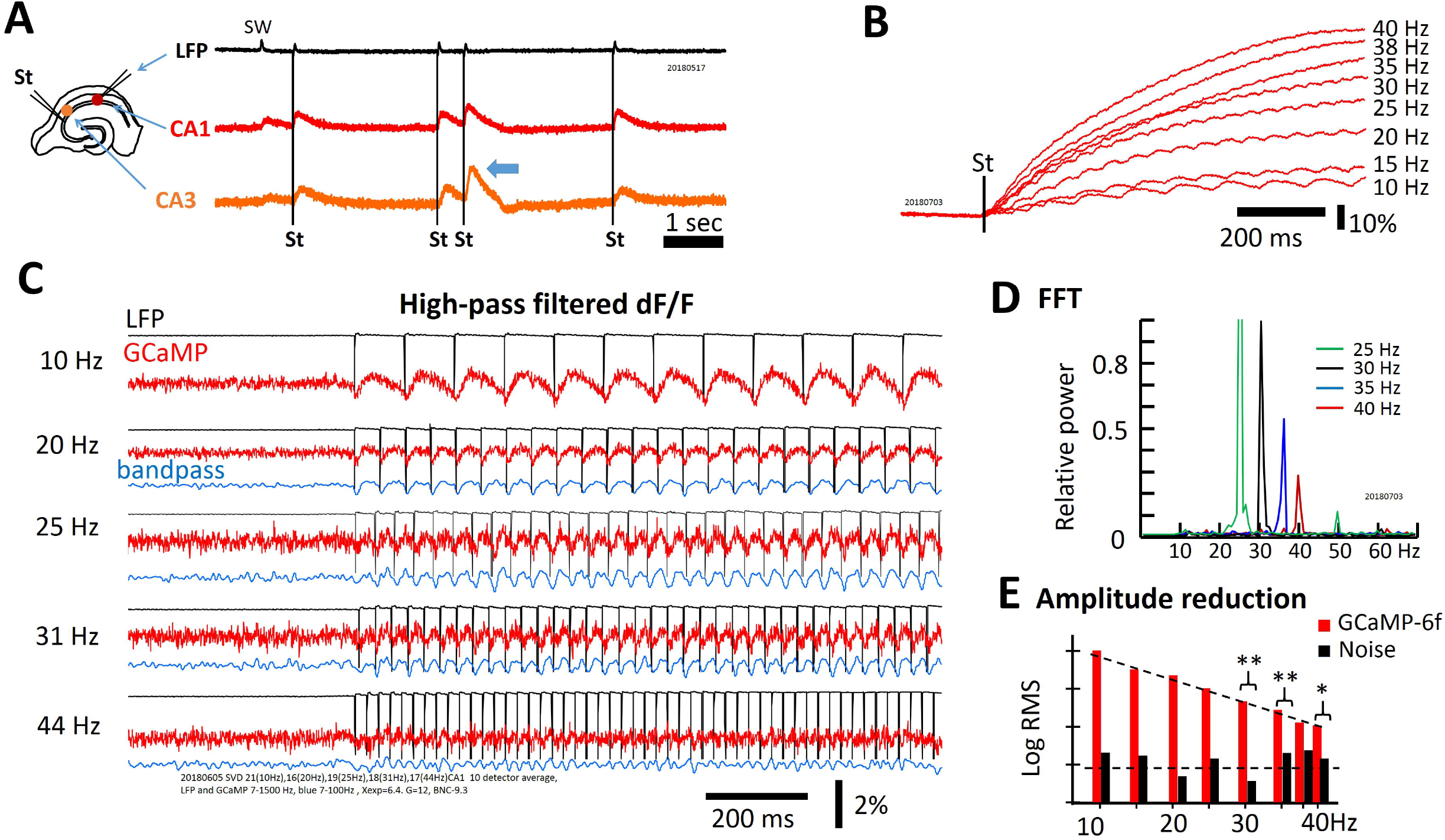
Frequency response property of GCaMP signals. **A.** GCaMP signals in response to mild electric stimuli. Left: The arrangement for the stimulation and recording. Stimuli were applied to CA3 and response was recorded in CA1 and CA3. The stimulation intensity was adjusted so that the evoked response had an amplitude similar to the SW amplitude in the LFP signals. Note that double stimuli induced larger response to the second shock, even with long inter-stimulus intervals of 500 ms (arrow head). **B.** High frequency stimuli caused a ramp accumulation of GCaMP signals. Note that the ramp signal was much large and slower than the response to individual stimuli. **C.** Evoked GCaMP signal to high frequency stimuli. High frequency component can be seen in raw signal (red traces, 5-200 Hz). Narrow band filtered (blue traces, 5-50Hz) improved signal-to-noise ratio. **D.** Power spectrum. Note that when the peak frequency power was normalized to 30 Hz, the signal power reduced to ~ 1/2 at 35 Hz and~1/4 at 40Hz. **E.** On a log scale the signal power quickly reduced in high frequency.

With a train of stimuli of identical intensity, the GCaMP signals summates, forming a much larger rising ramp than the response to individual stimulus (Figure 3B). The dF/F of individual stimulus was about 0.1%, while the ramp signal of the 40 Hz stimulation was ~100% (comparing Figure 3B and 3C). The ramp rising time was faster with higher stimulation frequency of identical stimulus intensity (Figure 3B). The large ramp signal suggested accumulation of calcium in the cell; the time course of the intracellular accumulation might be limited by the clearance of intracellular calcium, while the individual response might be related to the calcium influx.

The rising time of ramp signal was much slower than the rising time of individual response, so that the ramp signal can be removed by digital high-pass filter without significantly distorting the response to individual stimulus (Figure 3C). The one-to-one relationship between stimulus and GCaMP signals maintained up to 40 Hz of the stimulation frequency. Responses to <30 Hz stimuli were clearly seen in raw signals (Figure 3C). 30-40 Hz signals were distinguishable with band pass filtering (10 to 55Hz, blue traces in Figure 3C). Response to the 40 Hz stimuli was further verified with fast Fourier transform (FFT, Figure 3D). While the 40 Hz FFT peak was smaller compared to that of lower frequencies, the peak was clearly distinct from the background noise.

The power of population GCaMP signal decreased exponentially with frequency (Figure 3E), suggesting that the time delay of the GCaMP-6f limits the frequency response property.

### 3.6 Amplitude Discrepancy between GCaMP and LFP Signals

Exceptionally large GCaMP signals were occasionally seen during some population events, while the LFP signals of the same events were relatively small. Spontaneous interictal events has an amplitude of 17 + 12% (dF/F, N=15), in one typical preparation shown in Figure 4, about 20 times larger than the SW signals in the same tissue (dF/F 0.7+0.2%, N=455). In contrast the LFP signals of the SW and interictal spikes had similar amplitude albeit the polarity was reversed and more extracellular spiking were picked up by the LFP electrode (Figure 4A insert).

**Figure 4.**
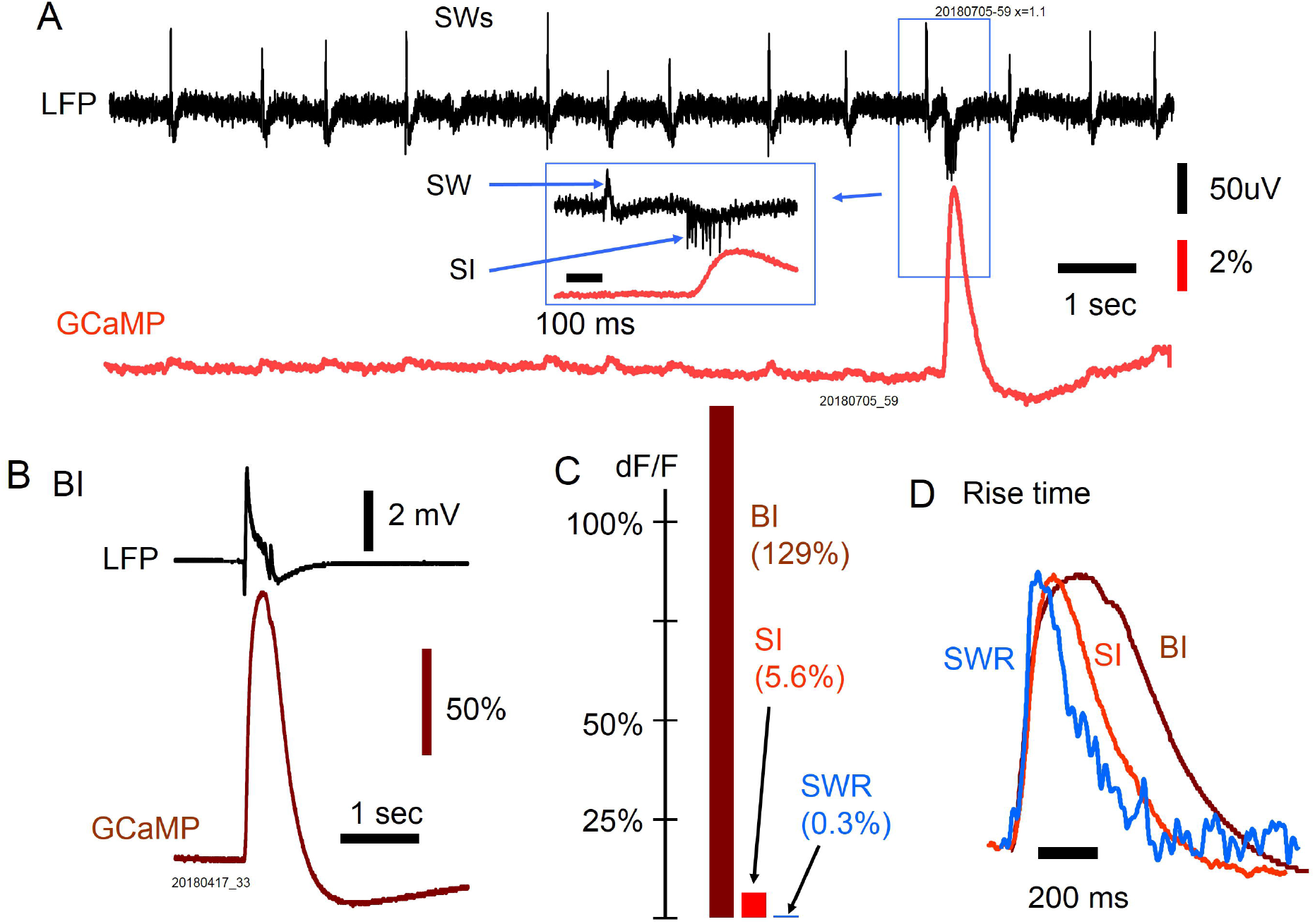
Disproportional between LFP and GCaMP signals. **A.** Spontaneous interictal event (blue box) associated with low LFP amplitude but unproportionally large GCaMP signals. Insertion: expanded time scale of the event, showing cellular spikes and the LFP peak of the interictal event. **B.** Interictal event induced by bath supply of 20 uM bicuculline. **C.** The GCaMP amplitude (dF/F) of different population events. **D.** LFP/GCaMP amplitude ratio of different events.

Spontaneous interictal events only occur occasionally in 2 out of 11 slices examined, and their occurrence rate was low. In one slice 1,847 SWs were recorded in 36 min of optical recording session, only 15 spontaneous interictal events were recorded. To further verifying the amplitude discrepancy, we examined bicuculline induced interictal events in the slices.

Bicuculline (20 μM in the perfusion solution) was used to induce spontaneous interictal-like spikes in 4 slices (N= 101 events recorded). The GCaMP signals of bicuculline induced interictal spikes were 200 - 500 times larger than the GCaMP of SW event, while the LFP signals was only ~20 times larger, demonstrating a large amplitude discrepancy. The dF/F of CCaMP signals expanded over a large dynamic range from SW to bicuculline induced interictal spikes, suggesting that GCaMP signals can be used to gauge the magnitude of population activity.

Surprisingly, the rising time for SWs and interictal spikes induced by the GABAa receptor antagonist was similar, despite the amplitude differ about 200 to 500-fold from SW to bicuculline induced interictal spike (Figure 4D). The rising time of all three events, defined as the time for the signal rising from 10% to 60% of the peak was about 40 ms. These suggest the onset time of the optical signal was limited by the GCaMP speed.

### 3.7 GCaMP Signals during Carbarchol Induced Theta Oscillations

Theta oscillations and related population events were used for further verifying the high frequency GCaMP signals and amplitude discrepancy with LFP recordings.

When the perfusant of the slice was switched from normal ACSF to ACSF containing 40 μm carbachol, the spontaneous SW disappeared and spontaneous theta oscillations (4-7Hz) emerged as recorded by the LFP electrode. Through time course transforming from SW to theta oscillations, large GCaMP signals and large discrepancy between the LFP and GCaMP signals were observed.

Figure 5A showed a representative recording through the transition from SW to carbachol induced theta oscillations. Before carbachol was introduced, slice in the control ACSF exhibit spontaneous SWs with LFP and GCaMP signals. These signals were used as a reference for the events later (Figure 5A trace 1, a section marked by the blue box is expanded in Figure 5B). Spontaneous SWs quickly disappeared when carbachol was introduced (within one min). Quickly, large fluctuations of GCaMP signals were observed, about 10 times larger compared to those of the SWs. Such large GCaMP fluorescence signal were correlated with small LFP signals, about 1/5 of that of the SW events. This again demonstrated amplitude discrepancy between LFP and GCaMP signals (Figure 5A, trace2, one section in blue box was expanded in Figure 5C). The optical signal fluctuation became larger and larger before theta oscillations emerged. Meanwhile the LFP electrode often picked up single unit spikes from neurons in the vicinity of the electrode tip, as a sign of elevated population excitability.

**Figure 5.**
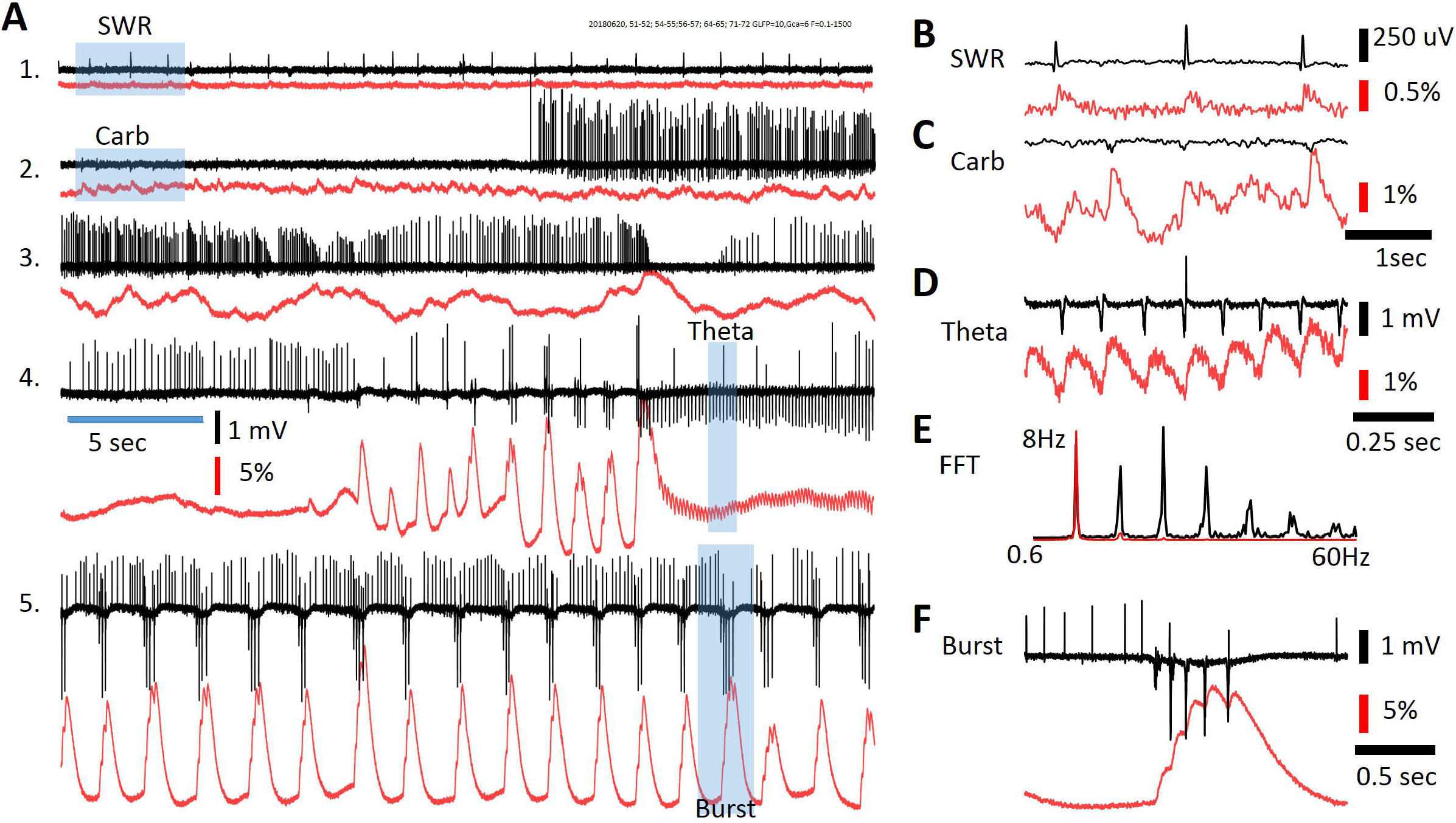
Comparison of LFP and GCaMP signals during carbarchol induced theta oscillations. **A1:** In normal ACSF, LFP and GCaMP signals show SW events. Black and red traces are for LFP and GCaMP signals respectively. Trace section in blue box expanded in **B. A2:** Just started (~1 min) perfusion with 40 uM carbarchol. Note that there are three changes in the signals: SW disappeared, irregular GCaMP signal fluctuations and elevated spontaneous firing from cells. GCaMP signal fluctuation and LFP peaks were correlated but amplitude disproportional. Trace section in blue box expanded in **C. A3:** later (~2 min) in carbarchol, increased spontaneous firing and larger GCaMP signal fluctuations. **A4:** Theta oscillations developed after large GCaMP signal fluctuations. Trace section in blue box expanded in **D. A5:** burst in theta activity. Trace section in blue box expanded in **F. E.** Power spectrum of the theta oscillation (in **A4**). Note that both LFP (black) and GCaMP (red) showed a primary peak of 8 Hz, LFP signals show more higher order harmonic peaks.

The GCaMP signal fluctuations paralleled changes in neuronal firing with bursts of spikes correlated with large increases of the GCaMP signal (Figure 5A, trace 3). Such correlation became more and more obvious, until an episode of theta oscillations occurred (Figure 5A, trace 4, a section in blue box was expanded in Figure 5 D). In this preparation the theta oscillation was about 8 Hz (Figure 5E), and the 8 Hz GCaMP signals were clearly seen (Figure 5D). Later, bursts of theta waves occur in the slice, each burst was associated with a large peak in the GCaMP signal, and a fast component correlated with each peak in the LFP.

During these events the discrepancy between GCaMP and LFP signals was large. In LFP signals the SW had similar amplitude compare to the theta burst peaks; in the GCaMP signals, the latter was 500-800 times larger. Again, the rising time of SW and theta burst were similar.

### 3.8 Cellular Transient and Population Signals

Finally, we wanted to verify that the cellular calcium transients were large, local to the soma, and had longer duration as reported before (Miyawaki et al., 2014; Norimoto et al., 2018). In contrast, the population GCaMP signals in the same tissue were small, distributed in a large area, and with shorter duration.

In an experiment showing in Figure 6, the CA3 area was imaged (Figure 6A) and the laminar pyramidale in the imagine field was examined for cellular calcium transients (Figure 6 A red hexagon). Five detectors were chosen to show both SW population signals (Figure 6B) and cellular transients (Figure 6C). In normal ACSF the SW signals were seen in all five detectors (Figure 6 B) as well as in most of the detectors in the imaging area (one SW marked in the blue box in Figure 6B was displayed in all detectors in Figure 6A). Later 2 μM carbachol was added into the perfusion solution to promote cellular spiking. Under this condition large calcium transients were observed in all five detectors. Because the diode array had a low spatial resolution, these calcium transients could not be attributed to individual CA3 neurons. However, these large signals were localized to one or a few detectors; e.g., the signals on neighboring red, blue, and green detectors only showed small crosstalk (Figure 6C), suggesting that the source of the signals was highly localized to the soma or dendrites of distinct CA3 neurons. In contrast, the SW signals were distributed over the entire field of view.

**Figure 6.**
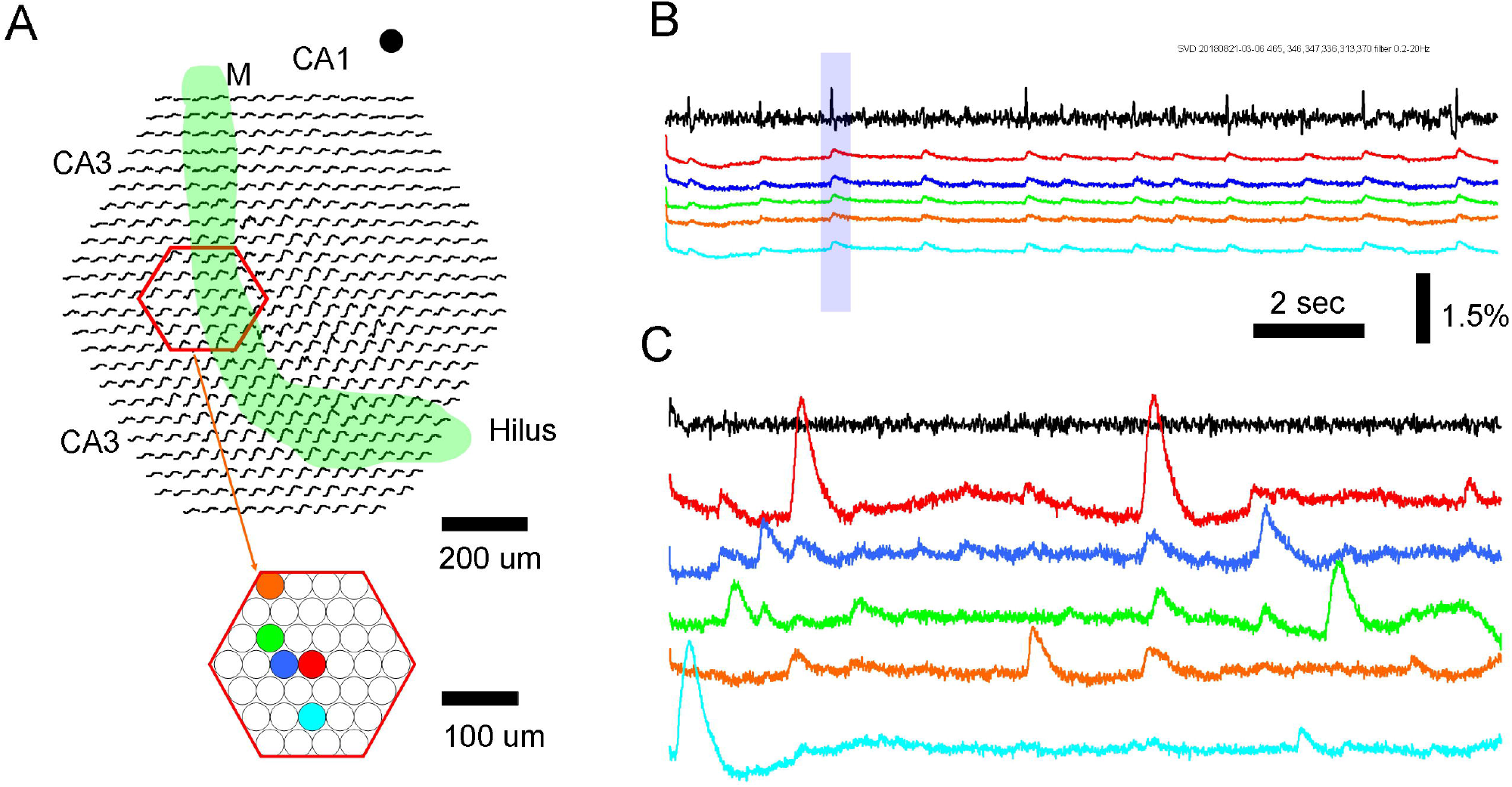
Population GCaMP signals of SW and localized cellular Ca transients. **A.** Schematic diagram of the imaging field. Green band marks the mossy fiber bundle. Red hexagon marks a group of optical detectors imaging the CA3 pyramidal cell layer. Color dots mark 5 individual optical detectors; signals on these detectors are displayed in the color traces in **B** and **C. B.** Traces of LFP (black) and GCaMP (colors) signals during SW events. LFP was recorded from a location in CA1, marked by the black dot on the top right of **A.** Color traces were from optical detectors (color dots in **A**). Blue box marked an example of SW event with optical recordings from the all detectors (black traces in the imaging field in **A**), demonstrating that a large fraction optical detectors around the mossy fiber bundle show SW signals. **C.** cellular Ca transients recorded from individual detectors in the CA3 pyramidal layer. Color traces are from detectors of the same color in **A.** The signals in B and C were from the same tissue, during normal ACSF (**B**) and 10 uM of Carbarchol (**C**). Note that the cellular Ca transients were localized, high amplitude and wide than the signals during SW.

The large and localized calcium transients also showed a duration of 1-2 seconds, consistent to the cellular calcium transients reported by other groups [e.g., (Dana et al., 2014; Miyawaki et al., 2014)]. The amplitude, spatial distribution and time course of the SW signals and the large-local calcium signals were all different, suggesting their sources were different, likely to be from population subthreshold event and cellular supra threshold events.

## 4 Discussion

Our results suggest that population calcium signal can be used as a new indicator for subthreshold and oscillatory activity in neuronal population. The range for detecting population signals spans 1000-fold, from ~0.1% to ~100% of dF/F. Similar to LFP recordings, it is sensitive to the population sum of subthreshold synaptic activities, such as SW, where majority of the CA1 neurons receive subthreshold synaptic potentials without firing action potentials (Hajos et al., 2013). Large LFP-GCaMP amplitude discrepancy were seen, suggesting that population calcium signals can be used for measuring population events invisible in the LFP signals, such as the large calcium fluctuations before carbachol induced theta oscillations emerged (Figure 5).

### 4.1 Time Delay

The time delay of the population GCaMP-6f signal was about 50 ms, which is slower than organic calcium indicators (e.g., Cal-520, Rhof-4, (Lock et al., 2015) reviewed by (Grienberger and Konnerth, 2012)]). The delay was likely to be caused by the response time of the protein. This limits the ability for measuring fast oscillations. Our results suggest that the speed of GCaMP-6f is sufficient for measuring oscillations below 20 Hz, while this can be pushed up to 30 to 40 Hz with offline signal processing. A better solution for detecting gamma oscillations would be faster calcium sensitive proteins [e.g., (Helassa et al., 2015; Helassa et al., 2016; Dana et al., 2018)].

### 4.2 Population Calcium Signal vs. Neuronal Calcium Transient

A major question raised by our results is whether the integration of CA1 cellular calcium transient would reproduce the population calcium signal seen during SW. Cellular calcium imaging during SW has been performed by the Ikegaya group from a large number of neurons with calcium transient in the CA1 area (Norimoto et al., 2012; Miyawaki et al., 2014; Norimoto et al., 2018). A careful analysis in these studies (Miyawaki et al., 2014) found that while 79% of the neurons showing calcium transients participate in SW events, each SW event only recruit ~4% of these neurons and each neuron only participates in ~5% of the SW events. Since ~70% of the neurons have calcium transients with and without SW, integrating the individual cellular calcium transient should not generate the population calcium signal as we observed in this report (see Figure 4 of (Miyawaki et al., 2014), also the supplement movie of (Norimoto et al., 2018)). Large fraction of uncorrelated cellular calcium transients would only contribute to the background fluorescent.

In contrast, the majority of CA1 neurons receives both excitatory and inhibitory synaptic inputs during every SW event (Hajos et al., 2013). The calcium influx in the presynaptic compartments of both excitatory and inhibitory neurons in the CA1 neuropil would contribute to the population GCaMP signal locked to the LFP, given that each CA3 neuron projects to 2/3 of the CA1 area and makes 30,000 to 60,000 of excitatory synapses onto CA1 neurons (Li et al., 1994; Wittner et al., 2007). The postsynaptic response of the CA1 neurons might also have active low threshold calcium channels [reviewed by (Catterall, 2000)] and contribute to the population calcium signal.

Cellular calcium transient reaches 30%-2000% of dF/F when recorded under a dark background with confocal or two photon microscopes. Under bright field fluorescent imaging the background is no longer dark so the fractional change would be greatly reduced. In our bright field fluorescent imaging the cellular calcium transient ranging from 2-5 times of the population GCaMP signals (Figure 6). The population signal of SW is a small intensity change over a brighter background (a light flux of ~ 10,000 photo electros/ms) which should saturate EMCCDs.

### 4.3 Amplitude Discrepancy

GCaMP-6f and LFP signals showed a striking amplitude discrepancy during some population events (Figures 4, 5). The unusually high amplitude in GCaMP-6f signal may reflect increased population firing rate in a duration of ~100 ms. The declining time of GCaMP-6f signal is about 200 ms (Chen et al., 2013), all calcium transients within the declining period should accumulate and integrate to the population signal. Another accumulation of the calcium signal is independent to the GCaMP declining time, but related to the summation of intracellular calcium. Individual neuron’s calcium transients last about 1 second in GCaMP-6f [Figure 6, see also (Chen et al., 2013; Dana et al., 2014) as well as measurements with organic calcium indicators (Miyawaki et al., 2014). The long duration of intracellular calcium transient might be related to the intracellular calcium buffering/elimination processes (Helmchen and Tank, 2015).

The accumulation of the population GCaMP signal can be clearly seen in Figure 3B, where repetitive stimuli caused a rising ramp and the ramp slope became deeper with higher stimulus frequencies. The ramp signal was much slower and larger compared to the signal induced by individual stimulus.

The accumulation of GCaMP offers a sensitive indicator for “the spiking density” in the population. Spiking density is referred to as increased firing rate on a temporal scale of 100 ms, which is different from the term “synchrony” as an increased firing rate on a millisecond temporal scale. High spiking density does not necessarily cause large LFP peaks, because sodium and potassium currents would cancel each other if the firing are not precisely synchronized. However, GCaMP-6f signal accumulates over ~100 ms time scale, offering an excellent indicator for increase spiking density. In contrast, on single cell level, the post synaptic potential and the GCaMP6 signal are much better correlated (Kupferschmidt and Lovinger, 2015). The large LFP-GCaMP discrepancy is likely to be a population sum of subthreshold phenomenon.

Large LFP-GCaMP amplitude discrepancy was seen in carbachol induced burst activity (Figure 5 A, trace 1 and 5). Asynchronized firing of large fraction of neurons during carbachol induced activity may explain the high amplitude GCaMP signals. In contrast, during SW only a few percent of neurons fire in a highly synchronized way, which would generate large LFP potential but small GCaMP signal.

### 4.4 SWR vs. Epileptic Events

There is an active debate whether in vitro SWRs reflect epileptic or other pathological events (Karlocai et al., 2014; Buzsaki, 2015). We demonstrated that the two events have large difference in GCaMP-6f signal amplitude. Spontaneous seizure events, while rare, can happen without changing the bath solution or the excitability of the slices. This provide a strong evidence that the in vitro SWRs are different from epileptiform events. The large calcium signals during epileptiform events are probably caused by large number of spikes in the neuronal population.

### 4.5 Sensitivity and Dynamic Range

We used diode array for our measurement, the limitation for the sensitivity seems to be the dark noise of the device. The intensity of excitation light needs to be high enough so the signal can be distinguished from the dark noise. In order to achieve ~30 min of optical recording time, we adjusted the excitation light so the dark noise was about 1/5 of the SW signals. (approximately 100 mW LED light output through a 10 x 0.3 NA objective, less than 1 mW/cm2 onto the tissue. Devices with lower dark noise (PMTs, EMCCD and cold CMOS camera) might get better sensitivity. Higher excitation power may also get better sensitivity with a trade in of optical recording time.

Dynamic range of the detecting device may also be a concern. An 8-bit of digitization resolution is needed for resolving the waveform of 0.1% dF/F and separating signal from the noise by digital filters. Another 8-12 bits are needed for 1000-fold change in the dF/F. These 16-20 bit dynamic range, 1000 frame/sec sampling speed would set a high bar for the imaging device.

Some of our results are compatible to a recent in vivo photometry study (Kupferschmidt et al., 2017). 1) high frequency stimuli generate a slow accumulative response and faster individual responses; 2) epileptiform events have high amplitude GCaMP6s signals. We were able to resolve fast signals up to 30 Hz on single trial, and able to record small signals (~0.1 dF/F) during hippocampus SWs. These probably because our diode array has higher dynamic range and signal-to-noise ratio than photomultiplier-based device. Further works are needed to verify if the high signal-to-noise ratio can be achieved under in vivo condition with optical fibers.

In conclusion, population GCaMP signals offer a sensitive way to measure asynchronized firing in a population event. It also allows no-contact recording and optical signals are not disturbed by the artifact of electrical stimulation. Sensitive and artifact free recordings are suitable for applications requiring recording at the same time of stimulation, e.g., interactions between EEG and transcranial repetitive AC or magnetic stimulation.

## 6 Author Contributions

PL and XG contributed equally. PL, HJ and JYW conducted the experiments. All authors participated in the data analysis and the composition of the manuscript.

